# The Epistasis Boundary: Linear vs. Nonlinear Genotype-Phenotype Relationships

**DOI:** 10.1101/503466

**Authors:** Serge Sverdlov, Elizabeth A. Thompson

## Abstract

Nonlinearity in the genotype-phenotype relationship can take the form of dominance, epistasis, or gene-environment interaction. The importance of nonlinearity, especially epistasis, in real complex traits is controversial. Network models in systems biology are typically highly nonlinear, yet the predictive power of linear quantitative genetic models is rarely improved by addition of nonlinear terms, and association studies detect few strong gene-gene interactions. We find that complex traits satisfying certain conditions can be well represented by a linear genetic model on an appropriate scale despite underlying biological complexity. Recent mathematical results in separability theory determine these conditions, which correspond to three biological criteria (Directional Consistency, Environmental Compensability, and Pathway Redundancy) together making up an Epistatic Boundary between systems suitable and unsuitable for linear modeling. For nonlinear traits, we introduce a classification of types of nonlinearity from a systems perspective, and use this to illustrate how upstream controlling genes, potentially more important to explaining biological function, can be intrinsically harder to detect by GWAS than their downstream controlled counterparts.

## 1. Introduction

The mathematical theory of separability, with roots in the work of Debreu (1960) and Luce and Tukey (1964) in the social sciences, can be applied to the problem of nonlinearity in the genotype-phenotype relationship.

The simple linear model that assigns an additive effect to each allele, e.g. ‘Replacing allele *a* with allele *A* always increases the specimen’s expected height by 0.3 cm,’ is understood to be an approximation. We replace the assumptions of this model with the less restrictive, more biologically plausible assumption of the directional consistency of substitution, e.g. ‘Replacing allele *a* with allele *A* always increases the specimen’s expected height.’ We find that traits meeting this and some additional assumptions can nonetheless be treated by linear model, when the scale on which the trait is measured is appropriately chosen. For such traits, which we call *separable*, apparent epistasis is an artefact of the choice of scale; these contrast with traits exhibiting *essential epistasis*, for which nonlinearity persists regardless of choice of scale, and making assumptions of linearity, e.g. in a GWAS context, can lead to directionally misleading results.

In translating the assumptions of separability theory to the genetic context, we address specific phenomena of genetics, namely diploidy, epistasis among loci, and the distinction between discrete genetic and possibly continuous environmental contributions.

### 1.1. The Epistasis Debate

It is a commonplace that the relationship between genotype and phenotype is complex and nonlinear. Models where the value of a trait is built up from additive contributions from each gene are present in every are of quantitative, statistical, and evolutionary genetic theory, and are often effective. Are such models merely convenient approximations, always wrong but sometimes useful? We find that there are biologically plausible circumstances under which linear modeling is appropriate, so long as the trait is expressed on an appropriate natural scale. A trait that satisfies a specified set of conditions will have this property; conversely, if a trait cannot be effectively represented by a linear model in our sense, that is, it exhibits essential epistasis, it must violate one of these conditions. In this way we hope to address the debate about the relative importance of epistasis in modelling the genotype-phenotype relationship, by classifying traits into two groups, linearly separable and non-separable, divided by the conditions of the Epistasis Boundary.

This debate touches on a fundamental question in genetics: can the effects of genes be understood separately, or must we consider each gene within the full complexity of its overall general genetic and environmental context? The controversy can be traced to the Haldane (1964) defense of “Beanbag Genetics,” the classical quantitative genetic models with simplifying assumptions including additivity of genetic effects, against Mayr (1963). Haldane advocated tractable idealized models, and analysis of deviations from idealized assumptions, against Mayr’s insistence on biological realism and complex models. Recent supporters of the linear approximation (e.g. Hill et al., 2008; Mki-Tanila and Hill, 2014), point to the empirical effectiveness of additive models in natural/evolutionary and artificial/agricultural selection and prediction, the limited empirical evidence for strong interaction effects, and the high fraction of variance captured by additive terms within a regression. Advocates of an important role for epistatic genetic variance (e.g. Zuk et al., 2012), argue both from natural complexity of biological mechanisms and nonlinearities produced by constructed system biology models. The large body of methods for detecting statistical epistasis in human populations (Wei et al., 2014), both as individual gene-gene interactions and globally, produces few positive results compared with model organisms (Huang et al., 2012; Mackay, 2014).

Contemporary reviews addressing the controversy (Cordell, 2002; Phillips, 2008; de Visser et al., 2011) emphasize the variety of concepts described by term epistasis, starting with the classical difference between biological epistasis (Bateson et al., 1905), the modification of the effect of an allele by an allele at another locus, and statistical epistasis (Fisher, 1918), defined in terms of residuals after subtracting effects explained by linear model.

## 2. General Nonlinearity and the Combinatorial Approach

Epistasis is one form of nonlinearity in the genotype-phenotype relationship; the others are dominance and gene-environment interaction. Our approach is directly applicable to dominance, and more easily demonstrated in that simpler context. More surprisingly, we will find that the general model incorporating environmental effects with epistasis is more tractable than the epistasis-only model, whether or not dominance is considered.

The general genotype-phenotype relationship 
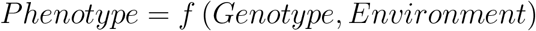
can be specialized by the assumption of no Gene-Environment interaction, or the additive separability of contributions from genotype and environment,

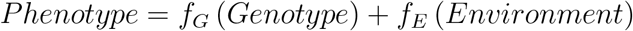

Further, the assumption of no epistasis implies additive separability over loci:

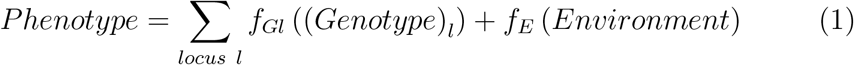

The assumption of no dominance deviation is equivalent to additive separability of the two diploid alleles within the same locus. Diploid symmetry implies that the functions mapping the two alleles at the same locus to their real-valued contributions are the same:

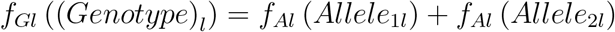

Such additive separability depends on the scale of measurement of the trait. If we observe a trait *T* without epistasis or dominance, measuring on a different scale, say ln *T* or *T*^3^, would violate additivity for any nontrivial choice of *T*, producing a biologically equivalent trait with epistasis or dominance. For a general scale transformation *S*, we cannot have both 1 and

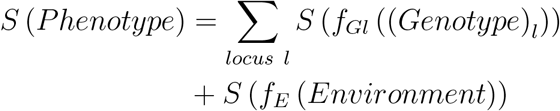

Thus some forms of nonlinearity are a mathematical artefact of the choice of scale, as opposed to an inherent biological property. Ideally, the question of whether epistasis or dominance exists as a biological phenomenon should be considered by a method invariant to monotonic transformation.

### 2.1. Natural Scale

This perspective leads us to consider ordinal, rather than quantitative, traits. By specifying traits as an ordering relation or ranking over genotypes, we can ask questions about traits without reference to the scale on which they are measured. In particular, can we choose a *natural scale* on which the trait is additive, that is, on which dominance and epistasis do not exist? Under what assumptions can such a scale be chosen, exactly or approximately?

Here we have formulated the natural scale problem in terms of biological epistasis, or nonlinearity within the genotype-phenotype relationship. In ??, we illustrate statistical epistasis from the population perspective, which manifests as a nonlinearity in the correlation function between relatives as a function of fraction of genome shared, under several models of genome sharing. We derive several generative models of gene-gene interaction that have this kind of nonlinearity as a signature. In statistical or population context, epistasis manifests as nonlinearity in the relationship between fraction of genome shared and trait correlation; in the biological or individual context, as nonlinearity in the genotype-phenotype relationship. In both cases, a transformation to the natural scale, if such a transformation is possible, eliminates epistasis. We will call such epistasis removable or *non-essential*, emphasizing the distinction between nonlinearity that is an artifact of the way we measure the trait, and the *essential* epistasis which is a biological phenomenon.

### 2.2. The Genotype Ordering Problem

Our method is based on the ordering of genotypes according to their representative phenotypic values. Such a ranking is not always possible to express unambiguously. A complete genotype-phenotype map incorporating all environmental inputs is deterministic, and each combination of genotype and environmental inputs corresponds to a single, and therefore rankable, phenotype. Likewise, we may treat a disease risk associated with a genotype, a single number, as a latent phenotype, and rank genotypes by disease risk. The problem arises with a classical quantitative trait like height, in which a genotype maps to a mean with a standard deviation associated with environmental variability. The straightforward solution would be to use the **genotypic value** defined as the average of the phenotype values for a given genotype, taken over potential realizations of the trait under environmental variability. However, this approach is problematic when we are considering nonlinear transformations of the measurement scale of the phenotype. There is no guarantee that order of genotypic values is preserved under such transformations. Table 1 illustrates this effect. We observe four phenotype values for each genotype; or, alternatively, the phenotype distribution given each genotype is discrete and consists of four equiprobable values. Three genotypes are ranked *AA ≻ AB ≻ BB* based on average trait value before transformation (upper table). The ordering is reversed to *BB ≻ AB ≻ AA* by logarithmic transformation (lower table).

**Table 1:** Counterexample to rank consistency of genotypic values under monotonic transformation.

|  | Phenotype Values |  |  |  | Average | Rank |
| --- | --- | --- | --- | --- | --- | --- |
| AA | 80 | 90 | 100 | 132 | 100.5 | 1 |
| AB | 80 | 100 | 100 | 120 | 100.0 | 2 |
| BB | 94 | 100 | 100 | 104 | 99.5 | 3 |
|  | Log Phenotype Values |  |  |  | Average | Rank |
| AA | 4.382 | 4.500 | 4.605 | 4.883 | 4.592 | 3 |
| AB | 4.382 | 4.605 | 4.605 | 4.787 | 4.595 | 2 |
| BB | 4.543 | 4.605 | 4.605 | 4.644 | 4.600 | 1 |

A sufficient condition for the ranking to be consistent is for the environmental variability to be homoscedastic, or additively independent. That is, on some scale, not necessarily the scale on which the trait is additive, the phenotype *P* can be decomposed into *P* = *G* + *E*, where *G* is the genotypic value and *E* is the environmental contribution, a random variable with a distribution that does not depend on *G*. Without this condition, genotypes cannot generally be ranked in a consistent scale-invariant way using average phenotypic value.

If, contrary to convention, we define genotypic value as the median, rather than mean, phenotypic value, we will not encounter this problem for continuous traits. The median would not be appropriate for discrete traits encoded as continuous variables, such as diseases with binary (0 or 1) status. For discrete traits, a genotypic value ranking can be associated with a risk or hazard rate, parameters which, though subject to nonlinear transformation with respect to e.g. time scale, maintain a consistent ranking.

## 3. Combinatorial Methods: Dominance in Small Diploid Systems

The choice of a natural scale is an assignment of real trait values to each genotype that maintains a given ordering of genotypes according to their representative phenotype values. Consider a single diploid locus with alleles *A,B,C,D*. The system is additive if the phenotypic value of genotype AB can be written as the sum of two allelic effects, e.g. *T*_AB_ = *A* + *B*. Several implications of additivity have a biological interpretation but are invariant to a monotonic transformation of the trait’s scale.

1. No overdominance (no cases such as *AA ≺ BB ≺ AB*)
2. No complete dominance (no *AA ≺ AB ≈ BB*)
3. Directional consistency of substitution (*CA ≺ CB* implies *DA ≺ DB*)

Each of these features is implied by additivity, and therefore necessary for the trait to be additive on some scale, but they are not sufficient. A transformation to additivity is not always possible; consider trait *T_ij_* for five alleles with ranking given by Table 2, which satisfies all three conditions. Suppose the trait was linearizable; then *A* + *D* = *B* + *C* and *C* + *D* = *A* + *E*; adding the two equations and subtracting *A* + *C*, we obtain 2*D* = B + *E* and therefore *f* (9) = *f* (10). We cannot find an additive solution with an increasing function *f*, though this particular contradiction does not prevent us from finding a non-decreasing solution, with *f* (*t*) increasing over *t* < 9 and *t* > 10 and constant for 9 ≤ *t* ≤ 10.

**Table 2:**
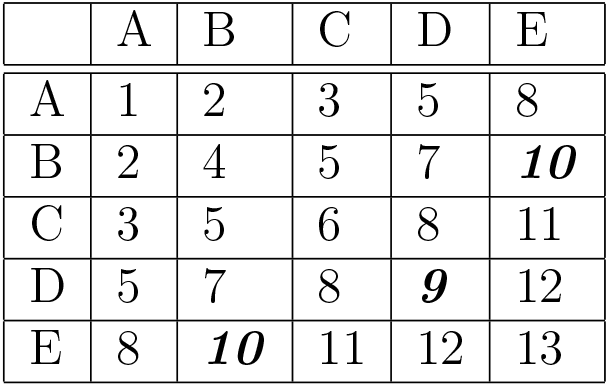
Trait with directional consistency of substitution, which provably cannot be transformed to additivity.

|  | A | B | C | D | E |
| --- | --- | --- | --- | --- | --- |
| A | 1 | 2 | 3 | 5 | 8 |
| B | 2 | 4 | 5 | 7 | **10** |
| C | 3 | 5 | 6 | 8 | 11 |
| D | 5 | 7 | 8 | **9** | 12 |
| E | 8 | **10** | 11 | 12 | 13 |

The ordering constraint on the genotypes of a single locus system imposed by non-overdominance or directional consistency of substitutionforms a directed acyclic graph (DAG). Any complete traversal of this DAG is an ordering of the nodes (genotypes) consistent with the ordering constraint, and therefore a trait architecture. For example, 
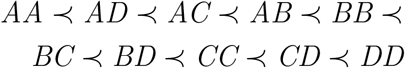
is an ordering that satisfies non-overdominance, but not directional consistency of substitution (e.g. *AC ≺ AB* but *BB ≺ BC*). The methods of Sverdlov and Thompson (in press) explicitly find transformations to the natural scale for separable or approximately separable traits. We applied these to all possible orderings for single locus systems satisfying the three criteria for up to 6 alleles. For systems up to 4 alleles all but two possible order architectures are solvable, that is, have a natural scale that respects the strict inequalities of the ordering constraints. We tabulated results for higher numbers of alleles as Table 3; the fraction of possible architectures that have an additive solution falls rapidly with number of alleles.

**Table 3:** Solvability of Dominance Systems for *n* = 1 *…* 6.

| $n$ | Total Architectures | Solvable Architectures |
| --- | --- | --- |
| 1 | 1 | 1 (100%) |
| 2 | 1 | 1 (100%) |
| 3 | 2 | 2 (100%) |
| 4 | 12 | 10 (83%) |
| 5 | 286 | 114 (39%) |
| 6 | 33592 | 2608 (7%) |

## 4. Discrete Separability Theory for Dominance and Epistasis

Separability, or additive utility, theory, concerns the conditions under which a weak order over tuples can be expressed as the order of sums of real-valued functions of the components of the tuples. The key early result in separability theory are due to Debreu (1960), in the context of a topological method with continuous variables applied to economic utility theory, and Luce and Tukey (1964) applied to measurement theory. Modern research efforts on the subject include those of Wakker (1989) and Gonzales (1996).

Our first interest is in additivity over finite, discrete sets. Theorems 1 and 2 (see Appendix A) follow Fishburn (1970), in the utility theory context, and we adapt the terminology to the genetic setting. An analogous but broader result, Theorem 4.1 in Fishburn (1970) has three forms (A, B, and C), which apply to different equivalence relations within the partial order (equality, equivalence, and indifference). The proof of this result can be based on, equivalently, the Theorem of the Alternative, Farkas’ Lemma, or the Separating Hyperplane Theorem.

Translating the notation to our setting, consider genotypes as multisets of alleles. This covers all cases of interest: single locus diploid as in the dominance model, and haploid or diploid multi-locus epistasis model. Each locus *i* = 1*…L* has a disjoint set of alleles *V_i_*. A diploid genotype over *L* loci contains 2*L* alleles, two from each *V_i_*, possibly the same allele repeated twice. The genotypic value function is given by a ranking over all realizable genotypes, without ties. The additivity question takes the form: under what circumstances can we assign real values to the alleles via a map *f*(), so that for all genotypes *X* and *Y,X ≻ Y* iff 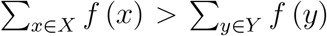. Applying Theorem 1, a linear solution *f*() exists if and only if there *does not* exist a paired list of genotypes *A*_1_ *…A_n_,B*_1_ *…B_n_* such that

1. the multiset unions of the two lists are the same
2. all *A_i_* ≽ *B_i_* and at least one *A_i_ ≻ B_i_*

We can state the principle in several different ways.

1. A trait possesses essential, non-removable epistasis if and only if, there is a finite population of individuals for which, by moving alleles between individuals without adding or removing any, the trait values of some individuals can be increased with none decreased.
2. In the non-epistatic case, if two populations have identical total gene content, one cannot dominate the other.
3. For an additive trait, reshuffling genotypes among individuals cannot increase the average trait value in a population.
4. For an additive trait, any population is Pareto optimal with respect to exchange of genes.

## 5. Continuous Separability Theory with Epistasis and Environmental Effects

A general genotype-phenotype relationship has the form 
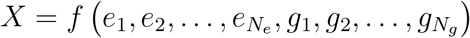
where *X* is a quantitative trait (e.g. plant weight), the *e_i_* are real valued environmental inputs (e.g. amounts of nutrients) and and *g_i_* encode the genotype. Theorems 3 and 4 (see Materials and Methods) describe the conditions under which a function of both real and discrete parameters is separable, i.e. transformable to the form 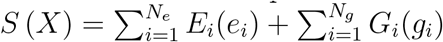. We translate these conditions into three principles interpretable in the context of our genetic and environmental variables. For expository purposes we will neglect mathematical detail and use a concrete example, referring to environmental inputs as nutrients, and to the trait as growth. The three principles are: Directional Consistency, Environmental Compensability, and Pathway Redundancy. Together these make up an Epistatic Boundary between separable traits, suitable to linear modeling, and non-separable traits, for which epistasis is present regardless of scale.

Under **Directional Consistency**, if a gene or nutrient is pro-growth, it’s always pro-growth in other environmental conditions and under any other genetic background. We consider directional consistency for substitutions of one gene or one environmental input at a time; that is, we change the value of one *g_i_* or *e_i_*. We do not require that a group substitution (e.g. turn off gene A and turn on gene B at the same time) have directionally consistent effect, though this more stringent condition is implied by separability.

**Environmental Compensability** is the ability to select a nutrient level that achieves any phenotype that is achievable through some other gene-environment combination. For example, if we remove a gene that contributes to growth, we can make up for that deficiency with additional nutrient. This is biologically plausible under midrange conditions of a complex system, but not in extreme conditions where a set of genetic substitutions changes the function of a pathway qualitatively rather than quantitatively.

**Pathway Redundancy** means we can achieve a trait value by multiple environmental means, or that environmental compensability applies to several environmental inputs. Three substitutable nutrient-type variables are sufficient; if there are only two, there are additional technical requirements about the tradeoff relationship between the two, and with only one the theorem provides no guarantees about separability.

Together, Environmental Compensability and Pathway Redundancy are the features of complex systems which are robust and redundant, and can smoothly make up for one input or capability by using up another resource. All three principles characterize maximization traits (e.g. speed, strength), traits for which it would be evolutionarily preferable to have even more if it didn’t involve tradeoffs of resources needed for other useful objectives, as opposed to traits with saturation or moderate-optimum properties.

The three principles (plus technical conditions) are sufficient for separability. Conversely, if the system is not separable, and there is non-removable epistasis that exists regardless of the scale, at least one of the three principles must be violated. A violation of directional consistency implies a logical switch, where a gene that is pro-growth in one set of circumstances becomes anti-growth in others. A violation of environmental compensability would occur if a genetic configuration leads to a trait outside the typical range of variability. An archetypal violation of pathway redundancy would be a bottleneck effect, where one input becomes a limiting reagent, and the system becomes highly sensitive to that input and indifferent to others. The hypothesis that individual genetic effects strong enough change the structure of the overall system will produce observable epistasis, associated with Crow (1990), corresponds to this kind of large scale change changing the critical path of the system. Analogously, the *decanalization* hypothesis (Gibson, 2009) holds that many genotype-environment combinations lead to normal phenotypes as biological systems stabilize a range of inputs, whereas disease phenotypes are attained either by extreme inputs or by breaking the system’s capability to stabilize, e.g by knocking out a hypothetical *robustness gene*.

## 6. Violations of Epistatic Boundary Principles and the Robustness Gene

Consider general ordinal traits, unconstrained by the epistatic boundary principles. The simplest such case is 2×2 tables, assigning rank values to the 2-locus, haploid system {*A*, *a*} × {*B*, *b*}. There are 4! = 24 possible orderings of the 4 cells of the 2 × 2 table. Without loss of generality, within-locus allelic types can be assigned to make *ab* the lowest ranked value, leaving 3! = 6 orderings. Likewise the labeling of *A* and *B* can ensure *Ab ≺ aB*. Only 3!*/*2 = 3 distinct rankings remain. For 2 loci, this problem has a decomposition into 3 cases; but with even 3 loci, the number of cases increase to 840. Table 4 illustrates this phenomenon. For a system with *n* (biallelic haploid) loci, there are 2^*n*^ genotypes, and therefore 2^*n*^! possible orders. Eliminating patterns equivalent up to symmetry, within-locus allelic types can be assigned to make *ab …* the lowest ranked value, and the labeling of *A, B, …* chosen to ensure *Ab · · · ≺ aB …*. This reduces this count of distinct patterns by a factor of 2*^n^n*!.

**Table 4:** Counts of general *n* × *n* epistatic order combinations.

| $n$ | Possible orders ($2^n!$) | Up to symmetry |
| --- | --- | --- |
| 1 | 2 | 1 |
| 2 | 24 | 3 |
| 3 | 40320 | 840 |
| 4 | $2.09 \times 10^{13}$ | $5.45 \times 10^{10}$ |
| 5 | $2.63 \times 10^{35}$ | $6.85 \times 10^{31}$ |
| 6 | $1.27 \times 10^{89}$ | $2.75 \times 10^{84}$ |
| 7 | $3.86 \times 10^{215}$ | $5.98 \times 10^{209}$ |

Table 5 illustrates the 3 patterns within the 2 locus system, named after the appearance of their orderings on a 2-by-2 table:

1. Pattern Z, directional consistency: *ab ≺ Ab ≺ aB ≺ AB*
2. Pattern, one-way inversion: *ab ≺ Ab ≺ AB ≺ aB*
3. Pattern X, two-way inversion: *ab ≺ AB ≺ Ab ≺ aB*

**Table 5:**
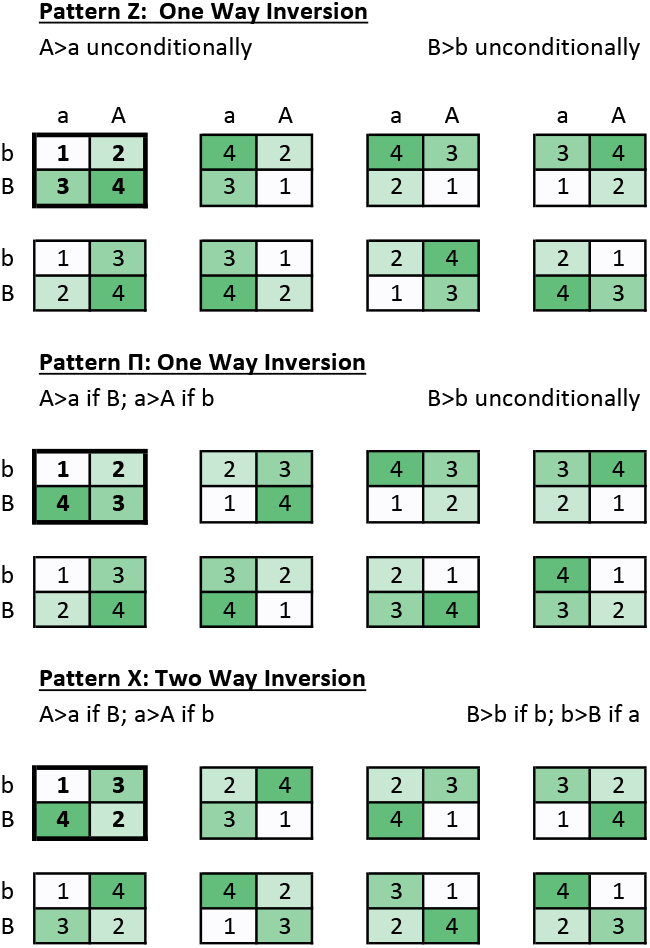
General 2 × 2 epistatic order combinations. 3 × 8 = 24 patterns reduce to 3 canonical patterns after fixing *ab* as minimum and *Ab ≺ Ba*.

|  |  |  |  |  |  |  |  |  |
|---|---|---|---|---|---|---|---|---|
|  | a | A | a | A | a | A | a | A |
| b | 1 | 2 | 4 | 2 | 4 | 3 | 3 | 4 |
| B | 3 | 4 | 3 | 1 | 2 | 1 | 1 | 2 |
| b | 1 | 3 | 3 | 1 | 2 | 4 | 2 | 1 |
| B | 2 | 4 | 4 | 2 | 1 | 3 | 4 | 3 |
| b | 1 | 2 | 2 | 3 | 4 | 3 | 3 | 4 |
| B | 4 | 3 | 1 | 4 | 1 | 2 | 2 | 1 |
| b | 1 | 3 | 3 | 2 | 2 | 1 | 4 | 1 |
| B | 2 | 4 | 4 | 1 | 3 | 4 | 3 | 2 |

Table 5: General $2 \times 2$ epistatic order combinations. $3 \times 8 = 24$ patterns reduce to 3 canonical patterns after fixing $ab$ as minimum and $Ab \prec Ba$ .
|  |  |  |  |  |  |  |  |  |
|---|---|---|---|---|---|---|---|---|
| b | 1 | 3 | 2 | 4 | 2 | 3 | 3 | 2 |
| B | 4 | 2 | 3 | 1 | 4 | 1 | 1 | 4 |
| b | 1 | 4 | 4 | 2 | 3 | 1 | 4 | 1 |
| B | 3 | 2 | 1 | 3 | 2 | 4 | 2 | 3 |

Pattern Z is the previously considered directionally consistent form. *a ≺ A* and *b ≺ B*, regardless of the state of the other gene. In Pattern X, switching either gene affects the other. The asymmetric character of pattern is the biologically interesting case, where one gene appears to control the action of another; this corresponds to the “robustness gene” hypothesis. One of the genes, *A*, can switch the direction of its estimated effect in the GWAS context, depending on the population frequency of *B*.

Consider a robustness gene *R* controlling a downstream client gene *C*. When the robustness gene is in the active state, the phenotype range is narrow, *f* (*Rc*) = 99 and *f* (*RC*) = 101; in the passive state, the phenotype range is wide, *f* (*rc*) = 80 and *f* (*rC*) = 120. Thus, the ordering is *rc ≺ Rc ≺ RC ≺ rC* and the pattern is ∏. Conditional on either *R* or *r, c ≺ C*; but *r ≺ R*|*c* and *R ≺ r*|*C*.

Note that even though biologically we would say *R* controls *C*, the direction of effect of *R* depends on *C*, not the other way around. In the GWAS setting, the effect of the *c → C* substitution is always positive. The effect of the *r → R* substitution changes sign depending on the population frequency of *C*. Thus, even though the *R* gene can be interpreted as more important, the *C* gene can be easier to detect by GWAS, and easier to replicate across different populations.

## 7. Discussion

Our method offers an explanation for the ‘unreasonable effectiveness’ of linear genetic models when the underlying biology is nonlinear, but satisfies our less binding conditions such as directional consistency.

### 7.1. Natural Scale

We introduced the concept of the *natural scale*, an exact or approximate tranformation of the quantitative phenotype on which Multiple theories within quantitative genetics are formulated with respect to additive models, and thus apply to the natural scale. The natural scale is the scale on which evolution operates in the sense of Fisher’s Fundamental Theorem of Natural Selection and the Breeder’s Equation. In the applied field of Genomic Selection, breeding value prediction and phenotype prediction methods work the same way under an exactly additive model, and differ when dominance is present. In the GWAS context, additivity has the implication that effects attributed to alleles do not differ due to allele frequencies; thus effects measured on an arbitrary scale should in principle vary from population to population, but not effects measured on the natural scale. Under our conditions, the same natural scale applies to environmental and genetic effects; with additional assumptions about many small independent effects, the central limit theorem implies normality of either the genetic or environmental contribution to the trait, on the same natural scale. Such a derivation of normality from theoretical conditions contrasts with methods such as Fusi et al. (2014), where a transformation to normality is used to pre-process traits for statistical analysis.

### 7.2. Epistasis Boundary

The Epistasis Boundary framework offers a conciliatory position in the debate about the importance of epistasis: epistasis can be neglected in favor of linear models for ‘well-behaved’ traits inside the boundary, and becomes relevant in more complex situations where the boundary’s constraints are violated. Figure 1 summarizes the decision criteria we have identified.

**Figure 1:**
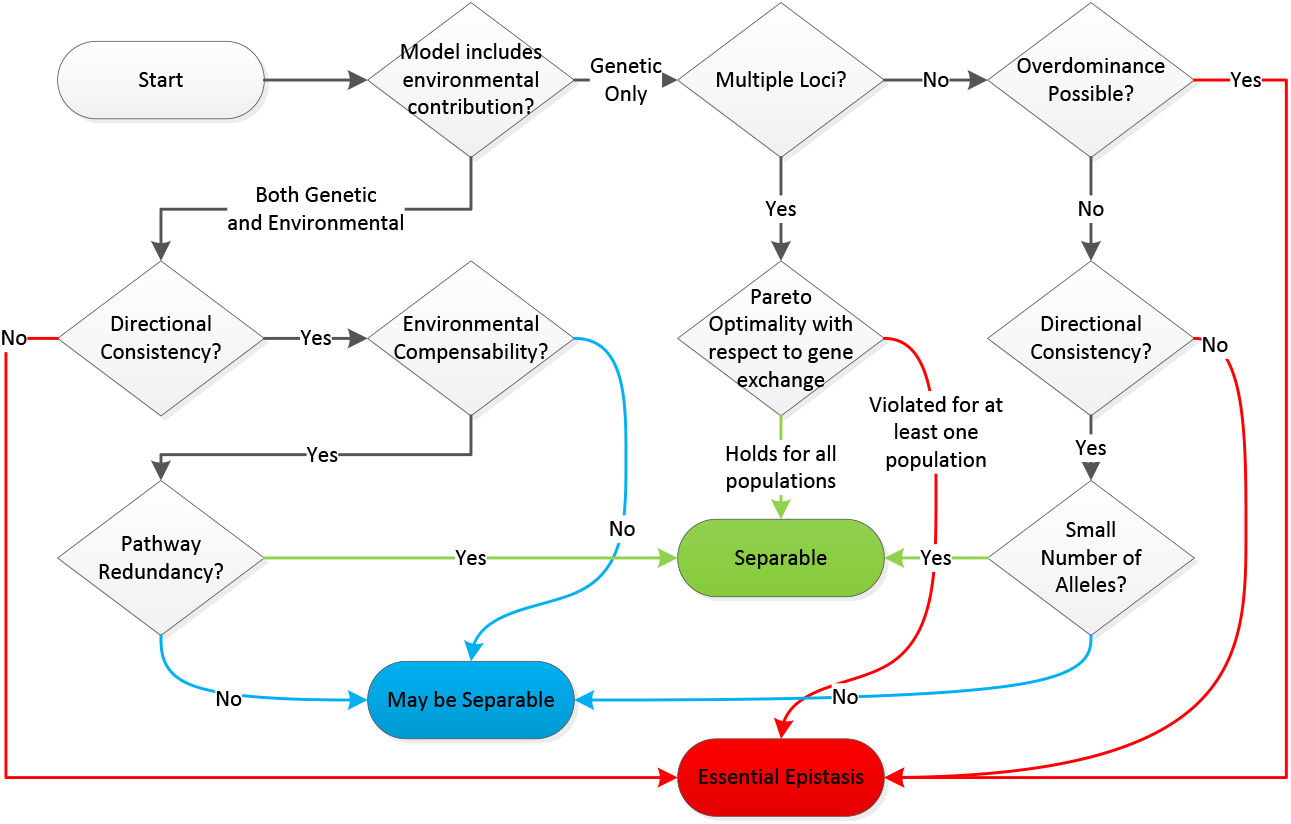
Decision Flowchart for Separability.

We can ensure that a trait is separable by meeting the combinatorial criteria identified for single locus systems, the population criteria for multiple locus systems, or our three principles for general systems with multiple loci and environmental contributions (Directional Consistency, Environmental Compensability, Pathway Redundancy). A separable trait can be represented by a linear model on an appropriately chosen scale. Violations of these criteria imply that a trait has essential epistasis that cannot be removed by a scale transformation. Such traits may are not suitable for linear modelling, and may exhibit phenomena such as the GWAS inversion we describe for the robustness gene, if linear models are inappropriately applied.

## 8. Acknowledgements

This research was supported in part by NIH grants R37 GM046255, T32 GM081062, and P01 GM099568.

## Appendix A. Definitions and Theorems

### Appendix A.1. Definitions

Following Fishburn (1970) and Fishburn (1992), let ≿ be a binary preference relation on a family *S* of subsets of a nonempty set *X*. Define the relations *A ≻ B* if *A* ≿ *B* and not *B* ≿ *A*; *A ∼ B* if *A* ≿ *B* and *B* ≿ *A*. The relation ≿ is **transitive** if for any *A, B, C ∈ S, A* ≿ *B* and *B* ≿ *C* together imply *A* ≿ *C*. It is **complete** if for any *A, B ∈ S*: *A* ≿ *B* or *B* ≿ *A*. It is a weak order if it is transitive and complete.

### Appendix A.2. Discrete Model

The pair (*S*, ≿) satisfies **model 1** if every *A* is finite there is a *u* : *X → R* such that for all *A, B ∈ S*,

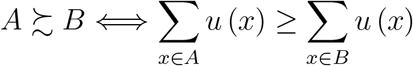

Let *A^′^* be the indicator function of *A ⊆ X*: *A^′^* (*x*) ∈ 1 if *x 2 A, A^′^* (*x*) = 0 otherwise. (*S*, ≿) satisfies finite cancellation if it is never true that there is a positive integer *m* and *A*_1_*, … A_m_, B*_1_*, … B_m_* in *S* such that

- 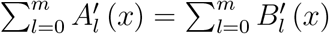 for all *x* ∈ *X*
- *A_i_* ≿ *B_i_* for all *i*
- *A_i_ ≻ B_i_* for some *i*

A collection of *∼* comparisons {*A_i_ ∼ B_i_, i* = 1*, … m*} satisfies linear independence if the *m* equations 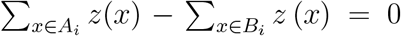 for each *i* = 1*, … m* are linearly independent (note that we are not defining *z*(*x*) but treating each *z*(*x*) for different *x* as a formal variable).

#### Theorem 1.

(from Fishburn (1992)) Suppose *S* is a finite collection of finite sets. Then (*S*, ≿) satisfies model 1 if (*S*, ≿) satisfies finite cancellation and ≿ is complete.

#### Theorem 2.

(from Fishburn (1992)) Suppose (*S*, ≿) satisfies model 1 under the initial conditions of Theorem 1, and |*X*| = *n*. Then *u* (*x*) is unique up to similarity transformations of the form *u^′^* (*x*) = *au* (*x*) + *b, a >* 0 if and only if some collection of *n* − 1 *∼* comparisons satisfies linear independence.

### Appendix A.3. Continuous Model

Following Gonzales (1996):

*Axiom 1*. (ordering) ≿ is a weak order on *X* (i.e. ≿ is complete and transitive).

*Axiom 2*. (independence) For any *i ∈* {*1, 2*, … *n*} and any *x, y ∈ X*, 
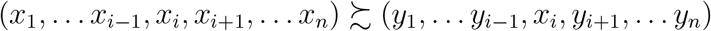
implies 

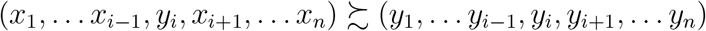

*Axiom 3*. (solvability with respect to the *i* th component) *x ∈ X, y_j_ ∈ X_j_* for all *j* ≠ *i* implies that there exists *z_i_ ∈ X_i_* such that

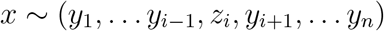

*Archimedean Axiom*. (Axiom 5 in Gonzales (1996)) Any strictly bounded standard sequence with respect to the first component is finite. For any set *N* of consecutive integers (positive or negative, finite or infinite), a set 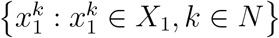 is a **standard sequence** with respect to the first component if not 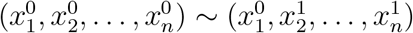 and for all *k* : *k, k*+1 *∈ N*, 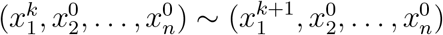.

#### Theorem 3.

(Theorem 1 in Gonzales (1996)) Assume that (*X*, ≿) is a weak order and that ≿ satisfies the independence axiom, as well as solvability with respect to the first two components, and that there exists an additive utility representing ≿_12_. Then there exists a unique additive utility representing ≿, that is, there exist real valued functions *u_i_* on *X_i_, i* = 1*, … n* such that for any 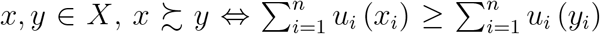 and these functions are unique up to similarity transformation.

#### Theorem 4.

(Corrolary 2 in Gonzales (1996)) Assume that (*X*, ≿) is a weak order and that ≿ satisfies the independence axiom, as well as solvability with respect to the first *three* components and the Archimedian axiom. Then there exists a unique additive utility representing ≿.

